# Dynamic fates of dietary antigen-specific T helper cells in a model of early life oral tolerance

**DOI:** 10.1101/2025.08.30.673288

**Authors:** Rachael D. FitzPatrick, Aleksandra M. Przydatek, Dominique M. Gatti, Natasha J. Norton, Jenna M. Lane, Brandon E. Moeller, Andrew N. Robinson, Lisa A. Reynolds

**Author notes:** Corresponding author: Dr. Lisa A. Reynolds. Mailing address: Department of Biochemistry and Microbiology, University of Victoria, PO Box 1700 STN CSC, Victoria, BC, Canada, V8W 2Y2.

## Abstract

Dietary antigens are first encountered in the gut during early life, when the immune system and microbiota are still maturing. In healthy individuals, oral tolerance develops towards dietary antigens: an active process which results in local and systemic immune unresponsiveness to antigens first encountered in the gut. Despite a wealth of research describing mechanisms contributing to oral tolerance in adult rodent models, questions remain about how the early life environment impacts oral tolerance development. We set out to characterize the fate(s) of CD4^+^ T cells during oral tolerance development using a robust early life mouse model, where controlled oral doses of dietary antigen are given directly to pups during the pre-weaning period. Orally administering 2mg of ovalbumin (OVA) daily during the third week of life was sufficient to confer oral tolerance to OVA in female and male C57BL/6 and BALB/c mice. Following early life oral OVA exposure, a large proportion of OVA-specific CD4^+^ T cells acquired a Th2 phenotype, alongside some OVA-specific Tregs. Following systemic challenges of OVA with an adjuvant, both OVA-specific Tregs and Th lineage-negative cells expressing anergy markers were detectable in pups given early life oral OVA, while OVA-specific Th2 cells were suppressed in comparison to pups who never received early life oral OVA. These data highlight the diverse fates of CD4^+^ T cells during early life oral tolerance development and maintenance, and we present a model to study oral tolerance during the early life period when dietary antigens are first encountered.

## Introduction

Oral tolerance is defined as the local and systemic immune unresponsiveness to antigens first encountered in the intestinal tract. The process of oral tolerance has been heavily studied, with initial hopes of its therapeutic application in the treatment of autoimmune and inflammatory diseases (1,2). Perhaps it is for this reason that the vast majority of studies describing the mechanisms that contribute to oral tolerance development have used adult rodent models, rather than focussing on how oral tolerance develops during the early life period when novel food antigens are initially encountered. The importance of early feeding of potential food allergens to infants for reducing the risk of later food allergy development is now appreciated (3). However, we are only just beginning to uncover how the capacity for and mechanisms supporting oral tolerance development may differ between the early life and adult setting (4–6).

It is well established that in adult mice, intestinal antigen-presenting cells present intestinal-derived antigens to naive CD4^+^ T cells in the gut-draining mesenteric lymph nodes (MLNs), a critical site for oral tolerance induction, and three fates of antigen-specific CD4^+^ T cells have been described: clonal deletion, anergy and/or differentiation into dietary antigen-specific Tregs (7). Dietary antigen-specific cells with features of anergy (Th lineage marker-negative, folate receptor 4 [FR4]^+^CD73^+^) that initially arise in response to oral antigen exposure can ultimately differentiate into Tregs to support oral tolerance development (8). These tolerogenic CD4^+^ T cell fates result in the inhibition of subsequent local and systemic adaptive immune responses, classically measured by an inhibition of dietary antigen-specific IgE responses after antigen challenges in inflammatory contexts (7).

However, studies of oral tolerance towards dietary antigens in adult rodents neglect to consider early life factors like an immature immune system, immature microbiome, and the consumption of breastmilk, all of which may impact oral tolerance development (4–6). Indeed, it has been known for many decades that neonatal mice (first week of life) do not have the same capacity to develop oral tolerance as adult mice (9,10), however, this capacity increases over the pre-weaning period (first three weeks of life) as levels of retinol increase (11), goblet cell-associated antigen cell passages form (12), and the numbers of antigen-presenting cells (Thetis cell type IV) peak in the MLNs (13). In early life oral tolerance models in rodents, the differentiation of dietary-antigen specific Tregs (12–14), the differentiation of dietary-antigen specific Th1 cells (11), and recently, the expansion of dietary-antigen specific CD4^+^ Th lineage-negative cells displaying anergy markers (15) have all been associated with efficient oral tolerance development.

Previous studies that have assessed dietary antigen-specific CD4^+^ T cell fates during oral tolerance development in the pre-weaning period have used models where young rodents are exposed to dietary antigens via breastmilk (11,12,14) or with dietary antigen also present in drinking water (13), making it difficult to control the exact antigen dose pups are consuming. While dietary antigen exposure via breastmilk mimics one physiologically relevant route that dietary antigen can be encountered during early life, we wanted to model the situation where novel dietary antigens are first being consumed directly by young animals, while they are also still drinking breastmilk. By administering dietary antigens directly to mouse pups, it is also possible to precisely control the dose of dietary antigen being consumed.

In the present study we demonstrate the use of a robust model of oral tolerance where mouse pups were directly orally administered physiologically relevant controlled doses of the model dietary antigen, ovalbumin (OVA), in their third week of life, prior to weaning. Using this model, we found that as expected, OVA-specific Foxp3^+^ Tregs expand following early life oral OVA exposure, but also that the majority of OVA-specific CD4^+^ T cells had acquired a Th2 phenotype (GATA3^+^Tbet^-^Foxp3^-^). However, following systemic challenges with OVA with an adjuvant later in life, mice that had received early life oral OVA had reduced frequencies of OVA-specific Th2 cells, and elevated frequencies of both OVA-specific Tregs and OVA-specific Th lineage-negative cells expressing anergy markers in comparison to pups who never received early life oral OVA. Together, these data characterize the fates of dietary antigen-specific CD4^+^ T cells during the development and maintenance of oral tolerance in early life and demonstrate the use of a robust model of early life oral tolerance during the pre-weaning period.

## Materials and methods

### Mice

All mouse experiments were performed at the University of Victoria (UVic), approved by the UVic Animal Care Committee and followed all guidelines set by the Canadian Council on Animal Care. Wild-type BALB/cJ (stock #000651), wild-type C57BL/6J (CD45.2; stock #000664), CD45.1 C57BL/6 (stock #002014), OT-II C57BL/6 (stock #004194), and ΔdblGATA C57BL/6 (stock #033551) mice were purchased from The Jackson Laboratory and subsequently bred in-house prior to use in experiments. Mice had access to food and water *ad libitum* and were group housed in individually ventilated cages under specific pathogen-free conditions. Wild-type and ΔdblGATA littermate male mice were generated by setting up breeding trios consisting of one hemizygous ΔdblGATA male with two females heterozygous for the ΔdblGATA mutation. Littermate male mice born to these heterozygous females were either of the wild-type or ΔdblGATA genotype. The dblGATA mutation is X-linked so it is not possible to generate wild-type and homozygous ΔdblGATA female littermate controls. Male littermates were continuously housed together after weaning until experimental endpoints.

### Oral tolerance to ovalbumin

Mice were orally gavaged with 2mg albumin from chicken egg white (ovalbumin; OVA, Grade V; Sigma-Aldrich #A5503) in 50μL of UltraPure™ water (Invitrogen) once daily for five consecutive days from days 16-20 of postnatal life. Control mice received oral gavages of 50μL UltraPure™ (Invitrogen) water. In some experiments mice were subsequently exposed to intraperitoneal injections of 100μg OVA and 1mg Imject™ Alum Adjuvant (Thermo Fisher Scientific) in a final volume of 100μL on days 22 and 36 of life. On day 43 of postnatal life, blood was collected from the brachial artery to quantify serum antibodies. All procedures were performed within 24 hours of the indicated age of mice.

### Quantification of antibodies by enzyme-linked immunosorbent assays (ELISA)

Serum samples were prepared by allowing blood to clot for 3-6 hours at 4°C then centrifuging blood at 9,600x*g* for 10 minutes. Serum was transferred to a new set of Eppendorf tubes and centrifuged for an additional 10 minutes at 9,600x*g*. Serum samples were stored at -20°C prior to assays. To quantify total IgG1 or IgE, MaxiSorp flat-bottom 96-well plates (Thermo Scientific™) were coated with either rat anti-mouse IgG1 (BD Biosciences; Clone A85-3) or rat anti-mouse IgE (BD Biosciences; Clone R35-72) diluted to 1:250 in 0.1M sodium carbonate (pH 9.5) overnight at 4°C. To quantify anti-OVA IgG1 plates were coated with 10μg/mL OVA (Grade V; Sigma-Aldrich #A5503) diluted in 0.1M sodium carbonate (pH 9.5) overnight at 4°C. All plates were blocked for 2 hours at 37°C with 2% bovine serum albumin (BSA; Sigma-Aldrich #A2153). A dilution series of each sample was incubated on plates overnight at 4°C alongside a standard curve of purified recombinant mouse IgG1 or IgE (BD Biosciences). Biotin rat anti-mouse IgG1 (BD Biosciences; Clone A85-1) or biotin rat anti-mouse IgE (BD Biosciences; Clone R35-118) were diluted to 1:1000 in blocking solution and incubated on plates for 1 hour at room temperature. Plates were then incubated with streptavidin-HRP (BD Biosciences) diluted to 1:1000 in blocking solution for 1 hour in the dark at room temperature prior to developing the ELISA with BD OptEIA™ TMB substrate (BD Biosciences). 1M hydrosulfuric acid was used for quenching. OVA-specific IgE was measured using a mouse anti-OVA IgE ELISA kit (Caymen Chemical Co.) following the manufacturer’s protocol. Samples were analyzed using a BioTek Epoch 2 microplate spectrophotometer (Agilent) and total IgG1, total IgE and anti-OVA IgE concentrations for each sample at a non-saturated dilution were interpolated from a standard curve. For OVA-specific IgG1 a standard curve was not generated, instead absorbance values were plotted at a non-saturated dilution.

### Ex vivo splenocyte culturing and cytokine quantification

Spleens were collected and cultured in sterile RPMI media (Sigma-Aldrich #R8758) containing 10% fetal bovine serum (FBS; Gibco), 0.1mg/mL streptomycin and 100 U/ml penicillin (Penicillin-Streptomycin solution; Sigma-Aldrich). Spleens were gently crushed through a 70μm cell strainer to achieve single cell suspensions, and red blood cells were lysed using ACK lysis buffer (Gibco). Cells were counted using a hemocytometer and 1×10^6^ cells were cultured in each well of a 96-well round bottom plate in the presence of media alone (unstimulated) or 10μg/mL OVA (Grade V; Sigma-Aldrich #A5503) or 1μg/mL anti-CD3 (BD Biosciences) as a positive control (data not shown) in a final volume of 200μL/well. Cells were cultured for 48 hours at 37°C and 5% CO_2_ and supernatants were collected and stored at -80°C prior to assays.

IL-5, IL-10 and IL-13 levels were measured in culture supernatants using the corresponding BD™ Cytometric Bead Array Mouse Flex Sets (BD Biosciences) according to the manufacturer’s protocol. Samples were analyzed using a CytoFLEX Flow Cytometer and CytExpert software (Beckman Coulter).

### Adoptive transfer of CD4^+^ T cells

The MLNs, inguinal lymph nodes and spleens were collected from donor male and female OT-II C57BL/6 (CD45.2) mice and gently crushed through a 70μm cell strainer into calcium- and magnesium-free phosphate buffered saline (PBS) solution containing 2% heat-inactivated FBS (Gibco) and 1mM EDTA (Invitrogen™). Red blood cells were lysed in spleen samples using ACK lysis buffer (Gibco). Splenocytes and lymph node cells were pooled from individual male mice or female mice, with male and female cells being processed separately. CD4^+^ T cells were isolated from each pooled sample using the EasySep™ Mouse CD4^+^ T cell isolation kit (STEMCELL Technologies) following the manufacturer’s protocol. Cells were counted using a hemocytometer prior to assessing the purity by flow cytometry and performing adoptive transfers. For purity testing, cells were stained with eBioscience™ Fixable Viability Dye eFluor™ 506 and Fc receptors were blocked with anti-mouse CD16/32 (2.4G2; BD Biosciences). Surface staining included APC-labeled anti-CD45.1 (A20; BD Biosciences), PE-labeled anti-CD45.2 (E50-2240; BD Bioscience), BV786-labeled anti-CD4 (RM4-5; BD Biosciences) and PeCy7-labeled anti-CD3 (145-2C11; BD Bioscience). OT-II cells were confirmed to be ≥90% pure for all experiments. Samples were analyzed using a CytoFLEX Flow Cytometer and CytExpert software (Beckman Coulter). 1×10^6^ CD4^+^ OT-II T cells were adoptively transferred in sterile PBS into sex-matched recipient CD45.1 C57BL/6 mice by retro-orbital injection into the venous sinus. OT-II C57BL/6 donor mice were 2-4 weeks old unless otherwise indicated.

### Cell isolations from tissues and flow cytometric analysis

MLNs and spleens were gently crushed through a 70μm cell strainer into PBS containing 0.5% BSA (Sigma-Aldrich #A2153). Red blood cells were lysed in spleen samples using ACK lysis buffer (Gibco). Resulting single cell suspensions from all tissues were counted using a hemocytometer prior to staining. Cells were stained with eBioscience™ Fixable Viability Dye eFluor™ 506 and Fc receptors were blocked with anti-mouse CD16/32 (2.4G2; BD Biosciences). Surface staining included: APC-labeled anti-CD45.1 (A20; BD Biosciences), PE-labeled anti-CD45.2 (E50-2240; BD Bioscience), BV786-labeled anti-CD4 (RM4-5; BD Biosciences), PeCy7-labeled anti-CD3 (145-2C11; BD Bioscience), APC-eFluor780-labeled CD44 (IM7;eBioscience), PeCy7-labeled FR4 (eBio12A5;eBioscience), and RB670-labeled CD73 (TY/11.8;BD Biosciences). For experiments where transcription factors were assessed, cells were permeabilized using an Invitrogen™ eBiocience™ Transcription Factor Staining Buffer Set following the manufacturer’s protocol, and transcription factor staining included: R718-labeled anti-T-bet (4B10; BD Biosciences), AlexaFluor488-labeled anti-GATA3 (L50-823; BD Biosciences) and V450-labeled anti-FoxP3 (MF23; BD Biosciences). For experiments where intracellular cytokines were assessed, cells were stimulated for 3.5 hours in the presence of 0.5 μg/ml phorbol 12-myristate 13-acetate (PMA) and 1 μg/ml ionomycin with 10 μg/ml brefeldin A included for the final 2.5 hours of culture at 37⁰C and 5% CO_2_. Following stimulation, cells were permeabilized using the BD Cytofix/Cytoperm™ Fixation/Permeabilization Kit (BD Biosciences) following the manufacturer’s protocol, and intracellular cytokine staining included: eFluor450-labeled anti-IL-13 (eBio13A; Invitrogen) and BV605-labeled anti-IL-4 (11B11; BD Biosciences). BD™ Liquid counting beads (BD Biosciences) were added to samples prior to acquiring data on a CytoFLEX flow cytometer (Beckman Coulter) to enable total cell counts according to manufacturer’s protocol. Data were analyzed using CytExpert software (Beckman Coulter).

### Statistical analyses

Datasets were first analyzed for normality using D’Agostino-Pearson omnibus normality and Shapiro-Wilk normality tests. Unpaired t-tests were used to assess differences between two normally distributed groups of data and Mann-Whitney tests were used to assess differences between two non-normally distributed groups of data. A Wilcoxon matched-pairs signed-rank test was used to assess differences between paired, non-normally distributed groups of data. GraphPad Prism software was used for statistical comparisons. A *p* value of ≤ 0.05 was considered statistically significant.

## Results

We set out to establish a model of oral tolerance development in the pre-weaning period where mouse pups are directly exposed to multiple precise physiologically relevant doses of the soluble dietary protein, ovalbumin (OVA), while they are still consuming breastmilk. We gave female and male mouse pups oral gavages of either control water or 2mg of OVA for five consecutive days during their third week of life (at days 16-20 old; **Figure 1A**). To assess whether these oral OVA exposures conferred oral tolerance towards subsequent systemic OVA challenges, we gave mice intraperitoneal injections of OVA with the IgE/IgG1-promoting adjuvant aluminum hydroxide (Alum) (16) and then assessed serum antibody levels (**Figure 1A**). C57BL/6 mice who were not exposed to oral OVA in early life developed OVA-specific IgE antibodies following systemic OVA/Alum challenges, while those mice who had received 2mg/day of oral OVA in early life exhibited a significant reduction in both OVA-specific and total IgE in their serum following systemic OVA/Alum challenges (**Figure 1B**). Similarly, OVA-specific IgG1 was produced in response to systemic OVA/Alum challenges in C57BL/6 mice who were not exposed to oral OVA in early life, but mice who had received 2mg of daily oral OVA in early life had significantly reduced OVA-specific and total serum IgG1 levels after systemic OVA/Alum challenges (**Figure 1C**). Together, these data indicate that 2mg of oral OVA administered for five consecutive days prior to weaning is sufficient for C57BL/6 mice to develop oral tolerance to OVA.

**Figure 1.**
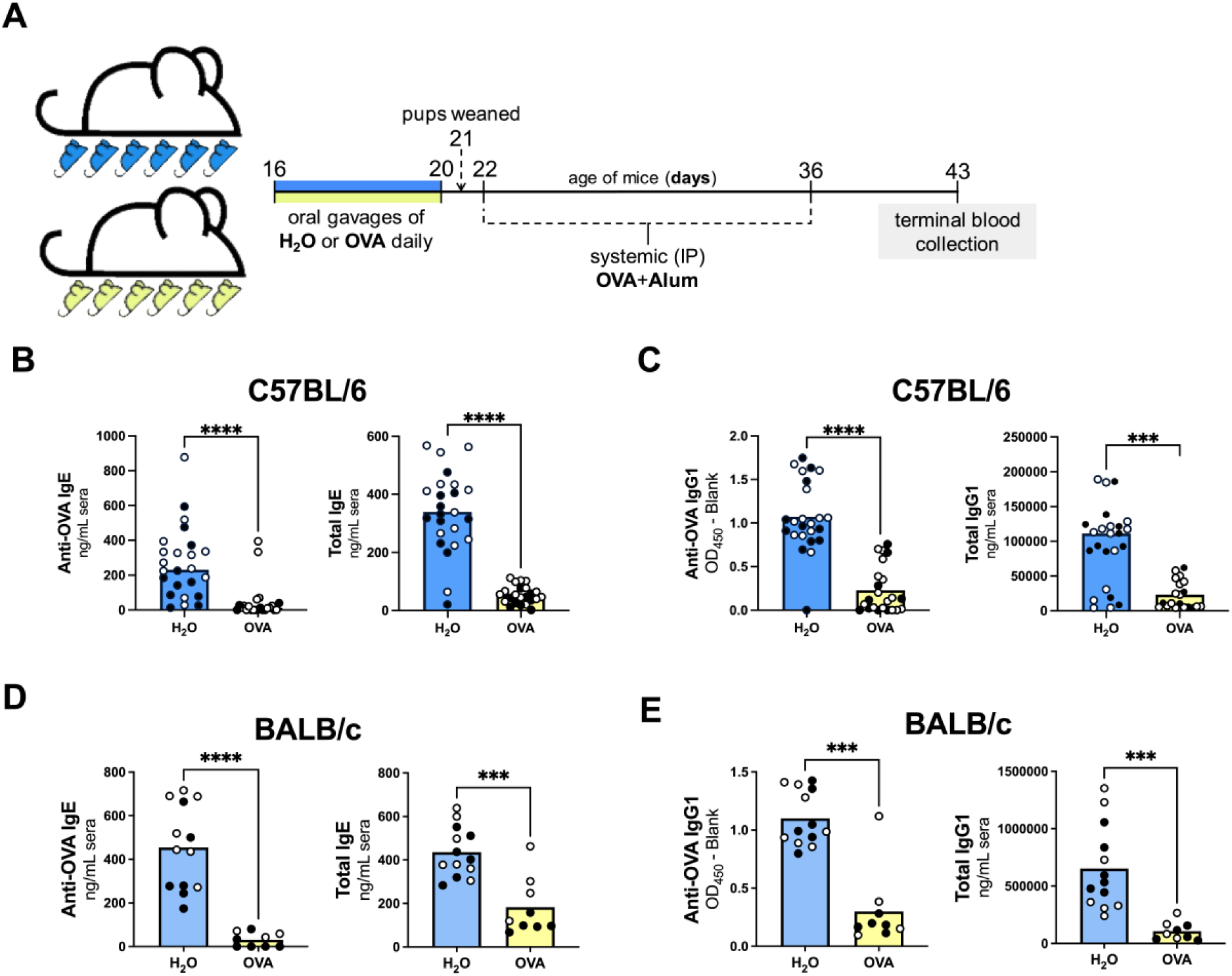
C57BL/6 and BALB/c mouse pups develop oral tolerance to 2mg daily oral OVA. **(A)** Experimental timeline: Male (●) and female (○) mouse pups were orally gavaged with H_2_O or 2mg of ovalbumin (OVA) in H_2_O on days 16-20 of life. All mice received systemic intraperitoneal (IP) challenges with 100μg OVA mixed with 1mg aluminum hydroxide (Alum) on days 22 and 36 of life. On day 43 of life blood was collected from each mouse to assess the development of oral tolerance. OVA-specific and total IgE **(B)** and IgG1 **(C)** were quantified by ELISAs in the serum of C57BL/6 mice exposed daily to H_2_O or 2mg OVA. OVA-specific and total IgE **(D)** and IgG1 **(E)** were quantified by ELISAs in the serum of BALB/c mice exposed daily to H_2_O or 2mg OVA. Column heights are at the mean and data were analyzed by an Unpaired t-test for normally distributed data. Column heights are at the median and data were analyzed by a Mann-Whitney test for non-normally distributed data. Data shown are pooled from 3 (B and C) or 2 (D and E) independent experiments with 4-9 mice per experimental group in each experiment. ***=p<0.001, ****=p<0.0001.

In some contexts, BALB/c mice exhibit exaggerated serum IgE and IgG1 responses when compared to C57BL/6 mice (17,18), and further, in a model where offspring were tolerized to an antigen via breastmilk, higher antigen doses were required to achieve tolerance in BALB/c pups compared to C57BL/6 mice (19). Therefore, we set out to confirm the efficacy of our early life oral tolerance model in BALB/c mouse pups. Consistent with our findings in C57BL/6 mice, we found that BALB/c mice exposed to 2mg daily oral OVA in early life had suppressed OVA-specific and total IgE and IgG1 (**Figure 1D+E**) levels following systemic OVA/Alum challenge compared to those who had not been exposed to oral OVA early in life, indicating these doses of early life OVA were sufficient to induce oral tolerance. Sex differences have been reported in IgE responses, whereby adult female mice develop stronger IgE responses following allergen exposure (20,21) and in humans, allergic males on average display elevated IgE levels compared to females (22). In our model, both female and male C57BL/6 (**Supplemental Figure 1A**) and BALB/c (**Supplemental Figure 1B**) mice were able develop oral tolerance after being exposed to 2mg daily of oral OVA early in life.

While antigen-specific antibody responses in sera are commonly used as a readout of oral tolerance development, we also wanted to assess systemic CD4^+^ T cell responsiveness to OVA following OVA/Alum challenge, in mice that had been exposed to early life oral OVA or not. To this end, we cultured splenocytes collected from individual mice after systemic OVA/Alum challenges in the presence or absence of exogenous OVA for 48 hours, prior to measuring cytokine levels in culture supernatants (**Figure 2A**). We found significantly reduced levels of the Th2-associated cytokines IL-5 and IL-13 in OVA-stimulated splenocyte cultures originating from mice who had received oral OVA in early life compared to mice who received control oral water in early life (**Figure 2B**), providing further evidence that the mice exposed to oral OVA had developed systemic tolerance to this model dietary antigen.

**Figure 2.**
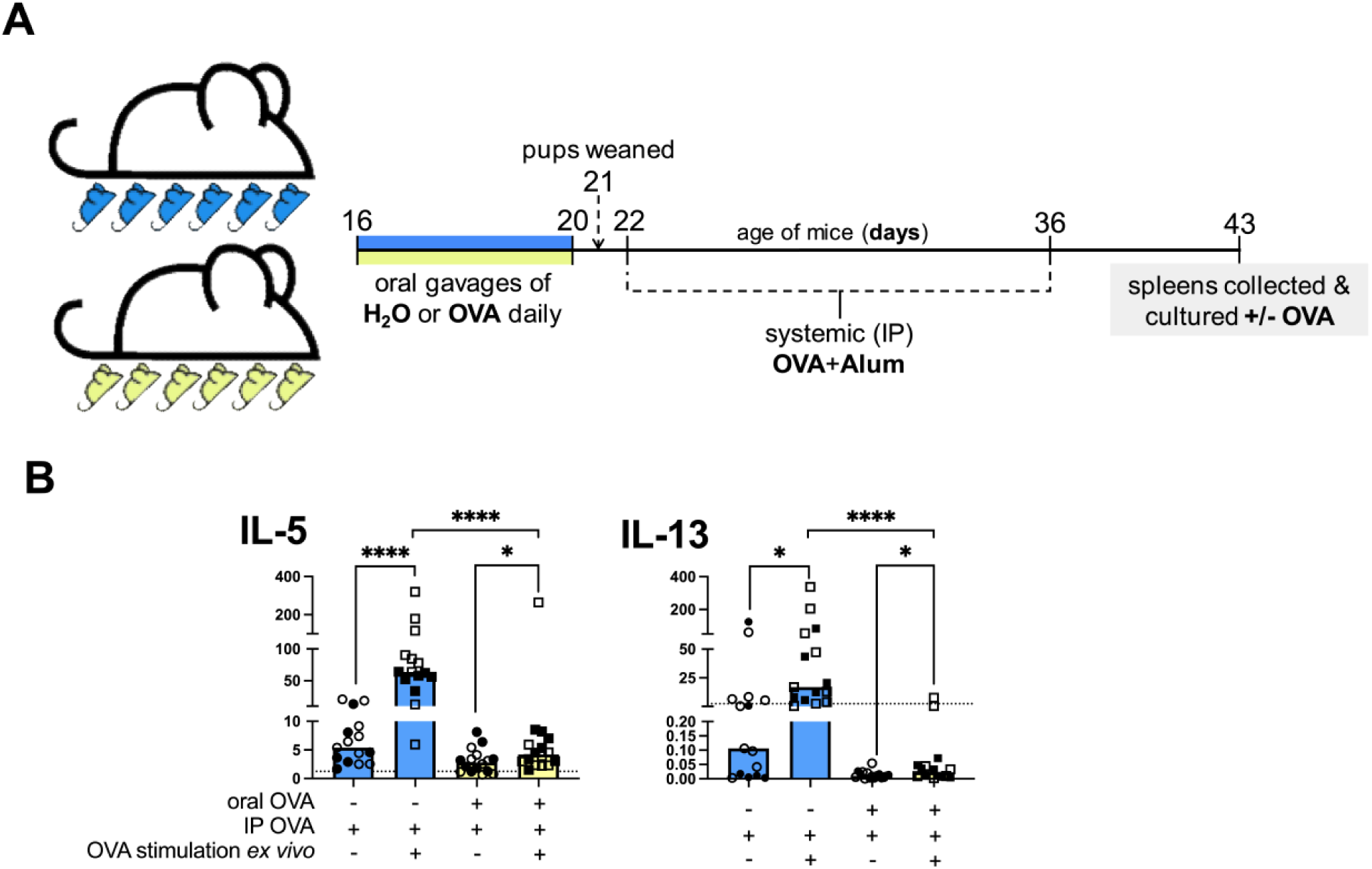
Early life oral OVA exposure leads to the suppression of OVA-specific Th2 cytokines following systemic challenge. (A) Experimental timeline: 16 day old male (●, ▪) and female (○, □) C57BL/6 mouse pups were orally gavaged with H_2_O or 2mg of ovalbumin (OVA) in H_2_O on days 16-20 of life. All mice received systemic intraperitoneal (IP) challenges with 100μg OVA mixed with 1mg aluminum hydroxide (Alum) on days 22 and 36 of life. On day 43 of life splenocytes were cultured in the presence or absence of 10μg/mL OVA for 48hrs and **(B)** IL-5 and IL-13 were quantified in culture supernatants using a cytometric bead array assay. Dotted lines indicate the lower limit of accurate quantification. Column heights are at the median and data were analyzed by a Wilcoxon matched-pairs signed rank test for paired experimental groups (unstimulated versus stimulated) and a Mann-Whitney test for unpaired experimental groups (oral H_2_O versus oral OVA). Data shown are pooled from 2 independent experiments with 7-9 mice per experimental group in each experiment. *=p<0.05, ****=p<0.0001.

In adult models of oral tolerance, eosinophils have been shown to contribute to tolerance development with eosinophil-deficient (ΔdblGATA) mice exhibiting less efficient induction of oral tolerance towards OVA when compared to wildtype mice (23,24). Eosinophils are present in the intestinal mucosa at comparable concentrations to that of adult mice within the first weeks of postnatal life (25,26), and so we set out to determine if eosinophils contribute to oral tolerance development in our early life model of oral tolerance. To best control for extrinsic factors, like exposure to differing microbes via the birth canal or breastmilk compositions in early life, we generated C57BL/6 wild-type and eosinophil-deficient (ΔdblGATA) male littermate pups. We found that, like in wild-type mice, ΔdblGATA mice given oral OVA in early life developed oral tolerance to OVA, with no differences in the levels of OVA-specific IgE, OVA-specific IgG1, or total IgE or IgG1 in tolerized mice between wild-type and ΔdblGATA mice (**Supplemental Figure 2A-C**). Our findings indicate that eosinophils do not contribute to the development of oral tolerance in this early life model.

Next, we set out to characterize the fates of OVA-specific CD4^+^ T cells after mice were orally exposed to OVA in early life. One possible mechanism contributing to oral tolerance is the clonal deletion of CD4^+^ T cells in the periphery, which has been demonstrated in adult OVA-specific T cell transgenic mice at higher dosages of oral antigen (27). To determine if clonal deletion was a driving mechanism of oral tolerance at our antigen dosage and during the early life time point, we transferred OT-II cells (CD45.2^+^) into 15 day old CD45.1^+^ C57BL/6 mouse pups prior to oral OVA exposure, allowing us to track the fate of OVA-specific T cells after mice were orally exposed to OVA in the context of an endogenous polyclonal T cell repertoire (**Figure 3A**). On day 22 of life, seven days after initial oral OVA exposure, we quantified OT-II cell (CD45.2^+^CD45.1^-^ CD4^+^CD3^+^) numbers in the MLN (**Figure 3B**) and spleen (**Figure 3C**), and did not detect any significant differences in the total number of OT-II cells between mice who had received oral OVA or those that had received oral H_2_O. These data suggest that clonal deletion of OVA-specific CD4^+^ T cells in the period immediately following oral OVA exposure does not drive the development of oral tolerance in this early life model.

**Figure 3.**
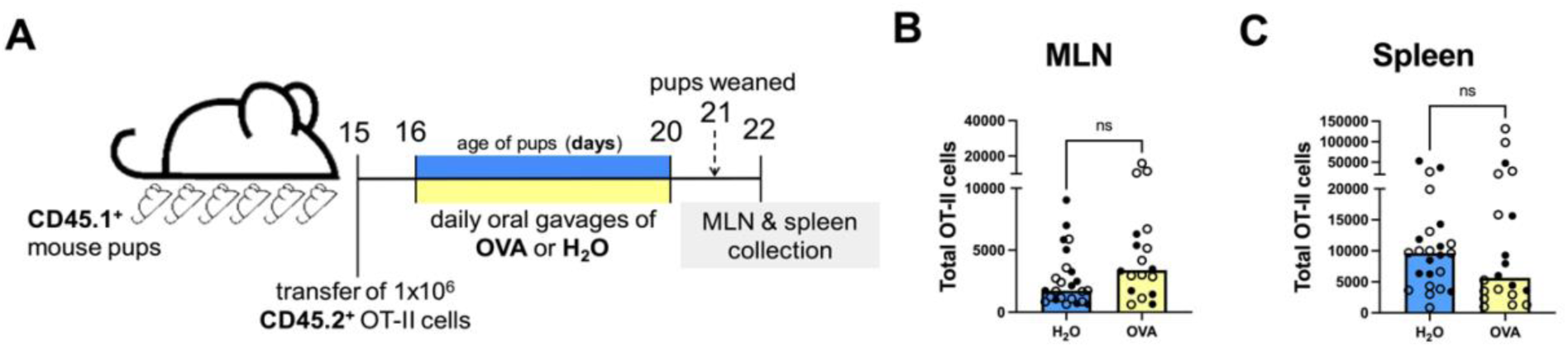
Deletion of OVA-specific CD4^+^ T cells does not drive oral tolerance to OVA in early life. **(A)** Experimental timeline: 15 day old male (●) and female (○) CD45.1 C57BL/6 mouse pups were injected with 1×10^6^ OT-II cells (CD45.2^+^) and then orally gavaged with 2mg of ovalbumin (OVA) in H_2_O or H_2_O alone on days 16-20 of life. On day 22 of life OT-II cells (CD45.2^+^CD45.1^-^CD4^+^CD3^+^ live singlets) were quantified in the **(B)** mesenteric lymph node (MLN)s and **(C)** spleens by flow cytometry. Column heights are at the median and data were analyzed by Mann-Whitney tests. Data shown are pooled from 4 independent experiments with 3-8 mice per experimental group in each experiment. ns = no statistical difference.

We went on to explore the phenotypes of adoptively transferred OT-II cells in the MLNs and spleen following early life oral OVA exposure (**Figure 4A**). Consistent with reports from adult rodent models and other early life models of oral tolerance (7,12–15), we found a significant increase in the frequency of OVA-specific Tregs (FoxP3^+^) in pups who were exposed to oral OVA compared to water-gavaged controls, in both the MLNs and spleen (**Figure 4B-C**). We next assessed whether we could detect any OVA-specific CD4^+^ T cells taking on a Th1 phenotype (T-bet^+^GATA3^-^FoxP3^-^) in response to oral OVA in early life, as it has previously been reported that the Th1-associated cytokine IFN-γ is required for BALB/c mouse pups to develop efficient oral tolerance to OVA that they have been exposed to through breastmilk (11). However, we detected essentially no OVA-specific Th1 cells in either the MLNs or the spleen of mice orally gavaged OVA or control water in early life (**Figure 4D** and data not shown). We next explored other potential Th fates of OT-II cells after early life oral OVA exposure and found an increased frequency of OVA-specific Th2 cells (GATA3^+^Tbet^-^FoxP3^-^) in both the MLNs and spleen of oral OVA-exposed mouse pups compared to water-gavaged controls (**Figure 4D-E**). To determine if OVA-specific CD4^+^ T cells also produced Th2-associated cytokines in oral OVA exposed pups, we stained adoptively transferred OT-II cells intracellularly for cytokines produced by Th2 cells (IL-4 and IL-13). Consistent with the higher frequency of GATA3-expressing OT-II cells in pups that were exposed to early life oral OVA (**Figure 4D-E**), we found a significantly higher frequency of IL-4- and IL-13-producing OT-II cells in the MLNs of mice exposed to early life oral OVA compared to control mice given water gavages, and a similar trend in the spleen which did not reach our threshold for statistical significance (**Figure 4F**). Supporting our findings of some OT-II cells acquiring both Treg and Th2 phenotypes following early life oral OVA exposure, we found that in a strictly endogenous setting where we gave wildtype C57BL/6 mice oral OVA or water in early life and then re-stimulated their splenocytes with OVA *ex vivo* (**Supplemental Figure 3A**), higher levels of Treg- and Th2-associated cytokines (IL-10, IL-5, IL-13) were produced in cultures from early life oral OVA-exposed pups compared to those from water-gavaged control mice (**Supplemental Figure 3B**).

**Figure 4.**
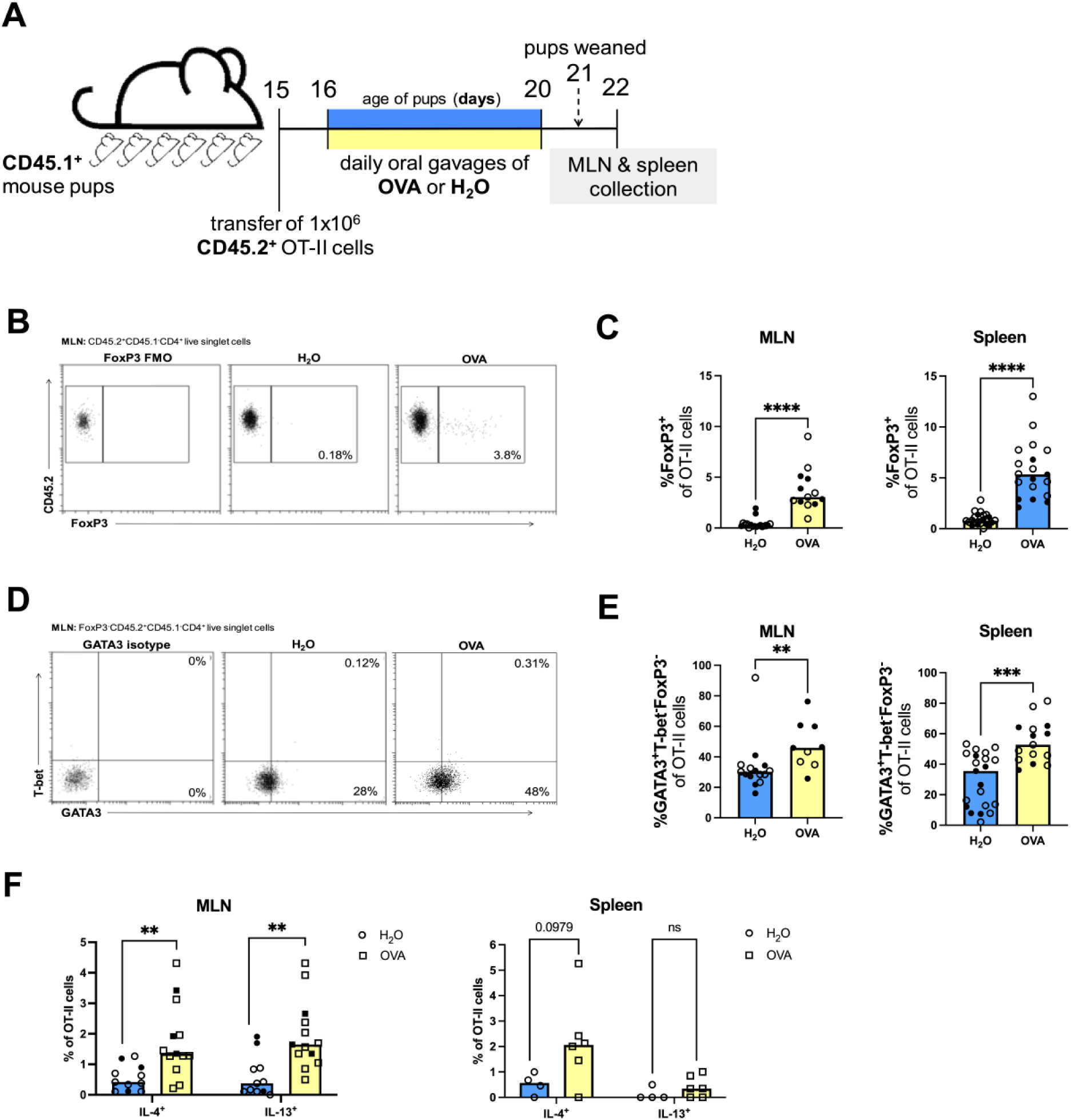
OVA-specific Tregs and Th2 cells develop after early life oral OVA exposure. **(A)** Experimental timeline: 15 day old male (●, ▪) and female (○, □) CD45.1^+^ C57BL/6 mouse pups were injected with 1×10^6^ OT-II cells and then orally gavaged with 2mg of ovalbumin (OVA) in H_2_O or H_2_O alone on days 16-20 of life. On day 22 of life the mesenteric lymph node (MLN)s and spleen were collected for flow cytometry. **(B)** Representative flow plots and **(C)** frequencies of Tregs (FoxP3^+^) among OT-II cells (CD45.2^+^CD45.1^-^CD4^+^CD3^+^ live singlets) in the MLNs and spleen. **(D)** Representative flow plots and **(E)** frequencies of Th2 cells (GATA3^+^T-bet^-^FoxP3^-^) among OT-II cells (CD45.2^+^CD45.1^-^CD4^+^CD3^+^ live singlets) in the MLNs and spleen. **(F)** The frequency of IL-4^+^ and IL-13^+^ cells among OT-II cells (CD45.2^+^CD45.1^-^CD4^+^ live singlets) in the MLNs and spleen. Column heights are at the median and data were analyzed by Mann-Whitney tests. Data shown are pooled from 3-4 (C), 2-3 (E), or 1-2 (F) independent experiments with 3-8 mice per experimental group in each experiment. **=p<0.01, ***=p<0.001, ****=p<0.0001, ns = no statistical difference, p values ≤0.1 are presented on graphs.

Many studies have suggested that in early life the adaptive immune system has a bias towards the development of Th2 responses (28). A contributing factor to this Th2 bias may be the higher proportion of recent thymic emigrant cells in younger mice (less than 6 weeks of age) within the peripheral T cell pool (29), which can have an increased propensity to differentiate into Th2 cells than mature naïve CD4^+^ T cells (30). Therefore, we set out to determine if the age of the donor OT-II mice may impact the differentiation of OT-II cells into Th2 cells after they were transferred into recipient pups that were then exposed to early life oral OVA (**Supplemental Figure 4A**). We did not find any increased intrinsic capacity for OT-II cells from young (3 week old) donors to differentiate into Th2 cells (GATA3^+^T-bet^-^FoxP3^-^) compared to OT-II cells from older (8 week old) donors after recipient pups were exposed to early life oral OVA: regardless of the age of OT-II donor mice, a sizeable proportion of OT-II cells had acquired a Th2 fate following oral OVA exposure (**Supplemental Figure 4B**).

We were surprised to find evidence for OVA-specific Th2 cell differentiation following early life oral OVA exposure (**Figure 4, Supplemental Figures 3+4**), given that we had demonstrated that early life oral OVA-exposed mice display oral tolerance to later challenge with systemic OVA/Alum as measured by the *suppression* of Th2-driven OVA-IgE, OVA-IgG1 and Th2-associated cytokines following systemic OVA/Alum challenge (**Figures 1+2**). Together, these data suggest that an immediate Th2 response to oral OVA exposure is later suppressed in response to systemic OVA/Alum challenges. To explore this, we characterized the Th fates of OT-II cells adoptively transferred into CD45.1^+^ C57BL/6 pups that were subsequently orally gavaged with either OVA or water and then challenged with systemic OVA/Alum (**Figure 5A**). Five days after systemic OVA/Alum challenge we quantified significantly fewer OT-II cells in the MLNs and spleen of mice orally gavaged with OVA in early life compared to water-gavaged controls (**Figure 5B**). When examining the frequencies of Th2 (GATA3^+^Tbet^-^Foxp3^-^) cells amongst antigen-experienced OT-II cells (CD44^+^CD45.2^+^CD45.1^-^CD4^+^; **Figure 5C**), we found lower frequencies of OVA-specific Th2 cells in the MLNs and spleen of mice exposed to oral OVA in early life compared to mice exposed to control water early in life (**Figure 5D**). In the spleens of these mice, we found a significant reduction in the frequencies of IL-4-producing OT-II cells in mice exposed to oral OVA in early life, but no statistical difference in the frequencies of IL-13-producing OT-II cells between mice exposed to oral OVA or water following systemic OVA/Alum challenge (**Figure 5E**). These data support a model where OVA-specific Th2 cells that expand after early life oral OVA exposure are later suppressed in response to systemic OVA/Alum challenge, resulting in suppressed systemic IgE and IgG1 responses and sustained oral tolerance in OVA-gavaged animals.

**Figure 5.**
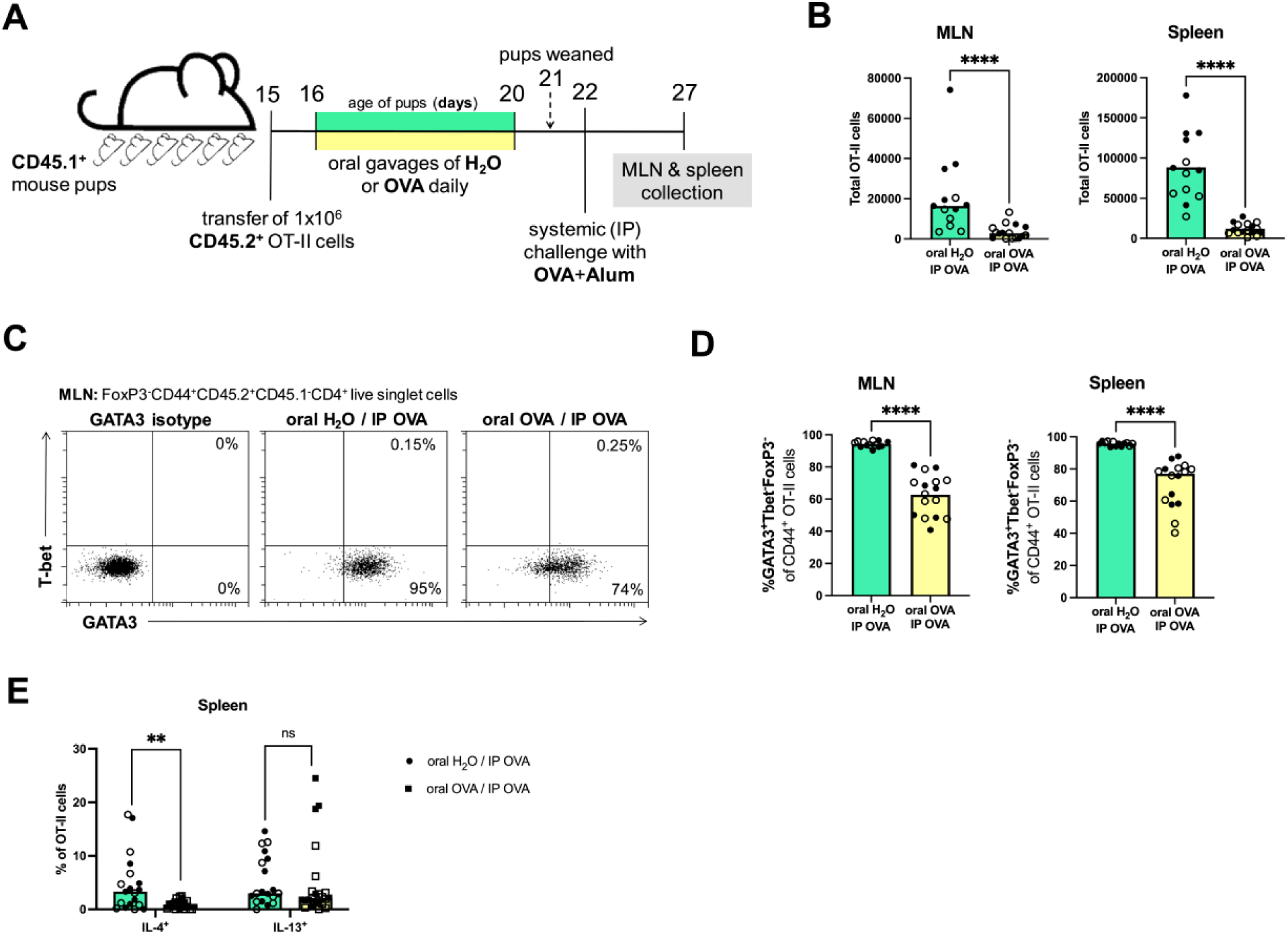
OVA-specific Th2 responses are suppressed following systemic OVA challenge in mice exposed to oral OVA. **(A)** Experimental timeline: 15 day old male (●, ▪) and female (○, □) CD45.1^+^ C57BL/6 mouse pups were injected with 1×10^6^ OT-II cells and then orally gavaged with 2mg of ovalbumin (OVA) in H_2_O or H_2_O alone on days 16-20 of life. All mice received systemic intraperitoneal (IP) challenges with 100μg OVA mixed with 1mg aluminum hydroxide (Alum) on day 22 of life. On day 27 of life the mesenteric lymph node (MLN)s and spleen were collected for flow cytometry. **(B)** Total number of OT-II cells (CD45.2^+^CD45.1^-^CD4^+^ live singlets) in the MLNs and spleen. **(C)** Representative flow plots and **(D)** frequencies of Th2 cells (GATA3^+^T-bet^-^ FoxP3^-^CD44^+^) among OT-II cells (CD45.2^+^CD45.1^-^CD4^+^ live singlets) in the MLNs and spleen. **(E)** The frequency of IL-4^+^ and IL-13^+^ cells among OT-II cells (CD45.2^+^CD45.1^-^CD3^+^CD4^+^ live singlets) in the MLNs and spleen. Column heights are at the mean and data were analyzed by an Unpaired t-test for normally distributed data. Column heights are at the median and data were analyzed by a Mann-Whitney test for non-normally distributed data. Data shown are pooled from 3 (B and D) or 4 (E) independent experiments with 4-8 mice per experimental group in each experiment. **=p<0.01, ****=p<0.0001, ns = no statistical difference.

We further explored the potential fate(s) of adoptively transferred OT-II cells following oral then systemic OVA exposure (**Figure 6A**). The expansion of dietary antigen-specific Tregs which can mediate oral tolerance towards dietary antigens has been well documented in adult models of oral tolerance (7), and in other early life models of oral tolerance (12–15). In addition, a recent study in adult mice suggests Th lineage-negative cells that display markers of anergy (FR4^+^CD73^+^) can contribute to oral tolerance development (8), and an elevated frequency of Th lineage-negative FR4^+^CD73^+^ cells in mice exposed to oral OVA at weaning has been associated with an increased capacity for oral tolerance (15). Consistent with these reports, after systemic OVA/Alum challenge we found elevated frequencies of both Tregs (FoxP3^+^) and Th lineage-negative cells expressing anergy markers (FR4^+^CD73^+^GATA3^-^T-bet^-^FoxP3^-^; **Figure 6B**) amongst antigen-experienced OT-II cells (CD44^+^CD45.2^+^CD45.1^-^CD4^+^) in the MLNs and spleen of mice exposed to oral OVA compared to mice who received control water in early life (**Figure 6C** and **D**). These data demonstrate that in the early life setting, both dietary antigen-specific Tregs and dietary antigen-specific Th lineage-negative anergic CD4^+^ T cells are associated with the development of oral tolerance.

**Figure 6.**
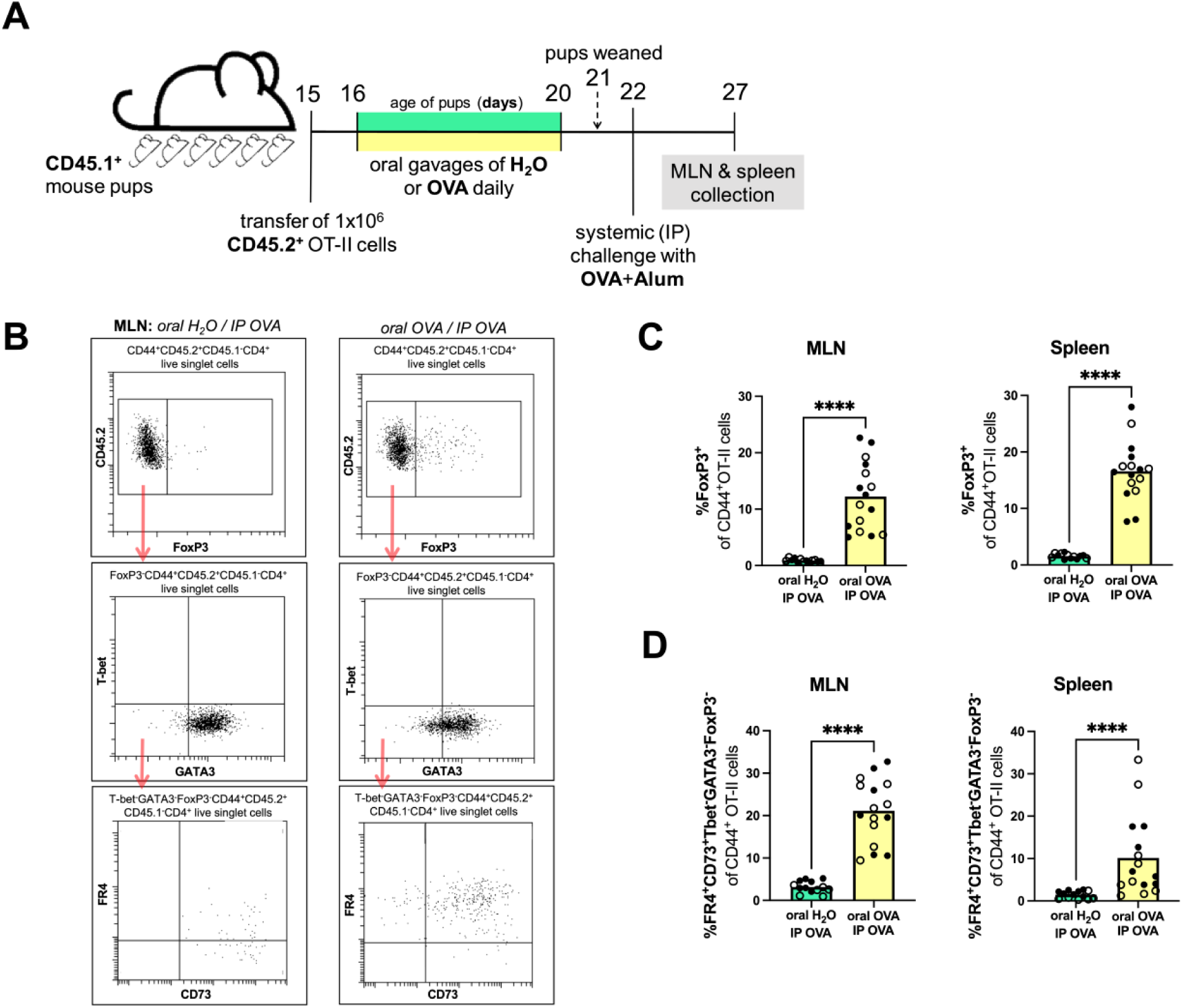
OVA-specific Tregs and OVA-specific Th lineage-negative FR4^+^CD73^+^ T cells are associated with early life oral tolerance development. **(A)** Experimental timeline: 15 day old male (●) and female (○) CD45.1^+^ C57BL/6 mouse pups were injected with 1×10^6^ OT-II cells and then orally gavaged with 2mg of ovalbumin (OVA) in H_2_O or H_2_O alone on days 16-20 of life. All mice received systemic intraperitoneal (IP) challenges with 100μg OVA mixed with 1mg aluminum hydroxide (Alum) on day 22 of life. On day 27 of life the mesenteric lymph node (MLN)s and spleen were collected for flow cytometry. **(B)** Flow cytometry gating strategies to identify: **(C)** frequencies of Tregs (FoxP3) among antigen-experienced OT-II cells (CD44^+^CD45.2^+^CD45.1^-^CD4^+^ live singlets) in the MLNs and spleen and **(D)** frequencies of Th lineage-negative anergic cells (FR4^+^CD73^+^GATA3^-^T-bet^-^FoxP3^-^) among antigen-experienced OT-II cells (CD44^+^CD45.2^+^CD45.1^-^CD4^+^ live singlets) in the MLNs and spleen. Column heights are at the mean and data were analyzed by an Unpaired t-test for normally distributed data. Column heights are at the median and data were analyzed by a Mann-Whitney test for non-normally distributed data. Data shown are pooled from 3 independent experiments with 4-6 mice per experimental group in each experiment. ****=p<0.0001.s

## Discussion

When oral tolerance fails to develop, food allergies can arise (7) and an increase in the prevalence of food allergies over the past several decades remains a concern. Historical feeding guidelines that urged parents to avoid feeding infants common food allergens have since been starkly reversed, to instead encourage early life feeding of potential food allergens (3). These guidelines shifted as a result of large randomised clinical trial studies, some of which demonstrated that early life consumption of certain potential food allergens was associated with reduced instances of allergies against those same foods later in life (3). Nevertheless, gaps remain in our understanding of how the immune system interacts with and develops tolerance to dietary antigens during the early life period. A deeper understanding of the mechanisms driving oral tolerance in the early life period may inform oral immunotherapeutic strategies to treat food allergy, or strategies to promote oral tolerance development in early life. In the present study, we set out to establish a robust mouse model of oral tolerance development where pups are directly exposed to a dietary antigen in their third week of life while they are still consuming breastmilk, which we used to investigate the fates of dietary-antigen specific T cells after initial early life oral antigen exposure and following subsequent systemic antigen challenges. We found that a dietary antigen-specific Th2 response initially develops alongside a dietary antigen-specific Treg response to oral dietary antigen exposure in early life, but this Th2 response is subsequently suppressed upon systemic challenge, and antigen-specific Tregs and Th lineage-negative cells expressing anergy markers expand in mice that exhibit oral tolerance.

A growing body of work is filling in the gaps in our understanding of factors that affect the capacity for oral tolerance development in the early life setting (4–6). Suckling mouse pups are not able to efficiently develop oral tolerance until their third week of postnatal life (9–12), with the maturation of their intestinal immune system and microbiota driven at least in part by components of breastmilk (12). These studies informed the timing we chose to directly introduce the model dietary antigen and common food allergen OVA to pups in our model: with daily oral gavages when pups were 16-20 days old. Our model aims to mimic a situation where an infant is first being exposed to novel dietary antigens in foods they are directly consuming, at a timepoint where they are also still consuming breastmilk. A benefit of this ‘direct-to-pup’ model is the ability to have precise control over the timing and dose of dietary antigen the pups are being exposed to. Many other groups studying oral tolerance development in early life have exposed pups to antigen via breastmilk (antigen delivered to dams) (4), which is another physiologically relevant route of first dietary antigen exposure, but lacks control of precise doses of antigen experienced by the pups. Like in adult models of oral tolerance, in early life oral tolerance models, dose of antigen experienced by pups influences the capacity for oral tolerance induction (4). Dietary antigen exposure via breastmilk, rather than direct-to-pup, can impact the process of oral tolerance development, as it can allow for the formation of maternal antibody-antigen complexes which pass to the infant via milk and are transcytosed across the infant’s intestinal epithelium via the neonatal Fc receptor, facilitating antigen-uptake by intestinal antigen presenting cells (4). Thus, the route, timing, and dose of dietary antigen exposure in early life are all factors that may affect the capacity for, and mechanisms of oral tolerance development in the early life period.

The mechanisms of dietary antigen uptake in the intestinal tract are unique at different life stages. In adult mice, dietary antigens are taken up in the small intestine and trafficked to small intestine-draining lymph nodes, which preferentially mediated tolerogenic responses compared to lymph nodes draining the distal intestine (31). In contrast, prior to weaning in mice, goblet cell-associated antigen passages located within the colonic epithelium are the major site of dietary antigen uptake, a mechanism driven by depleting levels of breastmilk-derived epidermal growth factor during this period (12). Very recent reports-mostly in adult mice-have identified subsets of variously named non-group 3 innate lymphoid cell RORγt^+^ intestinal antigen-presenting cells that facilitate the induction of dietary-antigen specific Tregs and mediate tolerance to dietary antigens (13,32–36). Notable among these reports is one describing how Thetis cell type IV antigen-presenting cells peak in abundance in gut-draining lymph nodes in the periweaning period, which promote the differentiation of dietary-antigen specific Tregs and oral tolerance development (13). Together, these studies further highlight potential differences in how oral tolerance develops in early life compared to in adulthood.

Using our pre-weaning model of oral tolerance development we set out to examine if, similar to reports in adult models (23,24), eosinophils contribute to oral tolerance development during the early life period. It is important to note that previous studies did not generate wild-type and eosinophil-deficient littermate controls for their experiments (23,24). Littermate controls are essential to control for extrinsic factors acting early in development that may impact lifelong immunity, for example, ingesting varying breastmilk compositions or exposure to different microbes via the birth canal (37,38). We compared oral tolerance development between male wild-type and eosinophil-deficient littermate mouse pups, born into the same cage from the same dam, and in our model found no impairment in oral tolerance in the absence of eosinophils. It is possible that the requirement for eosinophils in supporting oral tolerance development differs at different life stages, but it is also possible that extraneous variables arising due to a lack of littermate controls contributed to previous reports of a supporting role for eosinophils during adult oral tolerance development (23,24).

In our present study, we report that OVA-specific Th cell responses extend beyond the development of Tregs in response to oral OVA exposure in early life, as we also detected elevated OVA-specific Th2 cells in the MLNs and spleen of mouse pups immediately following oral OVA exposure. It is plausible the magnitude of this early antigen-specific Th2 response following oral OVA exposure is unique to the early life setting because the adaptive immune system is Th2-dominated during this period (28). We wondered if donor OT-II cells had an intrinsic bias towards Th2 differentiation if they were isolated from young, rather than adult, donor mice, however found that a substantial portion of OT-II cells transferred into early life oral OVA exposed recipients acquired a Th2 fate, regardless of the age of donor mice. We did not see evidence of the development of OVA-specific Th1 cells in response to early life oral OVA exposure in our model, in mice with a C57BL/6 background. This is in contrast to a previous report which showed OVA-specific Th1 cells developing which support early life oral tolerance development in BALB/c mice that were tolerized via OVA exposure in breastmilk (11); highlighting the likely impact of route of antigen exposure and/or antigen dose and/or genetic background on the mechanisms driving early life oral tolerance.

One recurrent theme between our present study, other early life oral tolerance models and in adult oral tolerance models is the differentiation of dietary antigen-specific Tregs following oral dietary antigen exposure (7,12–15). In an adult mouse setting, where dietary antigen-specific CD4^+^ T cell responses were tracked within a polyclonal T cell repertoire using tetramer-binding enrichment, diverse fates of dietary antigen-specific CD4^+^ T cells were reported following dietary antigen exposure, which included differentiation of Tregs, but also expansion of a population of Th lineage-negative cells expressing anergy markers (FR4^+^CD73^+^) (8). These authors went on to demonstrate that Th lineage-negative cells could ultimately develop into Tregs that contributed to oral tolerance development (8). In our early life oral tolerance model, we also saw expansion of OT-II cells that were Th lineage-negative FR4^+^CD73^+^, in mice that had received early life oral OVA, but not in mice that had received only oral water in early life. A recent study where mice were first exposed to oral OVA at the point of weaning demonstrated that exposure to antibodies in breastmilk early in life helped to limit inflammatory T cell responses to OVA at weaning, and intriguingly, the presence of antibodies in breastmilk was associated with increased numbers of Th lineage-negative FR4^+^CD73^+^ cells, but not Tregs, which suggests Th lineage-negative FR4^+^CD73^+^ cells may support oral tolerance development in the early life setting (15), like in adulthood (8). Whether Th lineage-negative FR4^+^CD73^+^ cells that develop after early life dietary antigen exposure also go on to acquire a Treg fate has yet to be explored.

In summary, we have demonstrated the use of a mouse model of oral tolerance development during a pre-weaning period of life when pups have the capacity to develop oral tolerance, are still consuming breastmilk, and where antigen is administered directly to pup. Using this model, we determined that eosinophils are not an essential cell type for oral tolerance development in early life. Further, we used this model to highlight the diverse and dynamic fates of dietary antigen-specific Th cells in response to initial early life oral antigen exposure, and during oral tolerance maintenance.

## Acknowledgements

We would like to thank the UVic Animal Care Staff for providing excellent care to the mice used for this project. We greatly acknowledge the animals used in this study and recognize their invaluable contribution to advancing scientific knowledge.

## Author contributions

R.D.F.: Conceptualization, Data curation, Formal analysis, Investigation, Methodology, Project administration, Supervision, Validation, Visualization, Writing- original draft, Writing- review & editing.

A.M.P.: Data curation, Formal analysis, Investigation, Writing- review & editing.

D.M.G.: Investigation, Writing- review & editing.

N.J.N.: Investigation, Writing- review & editing.

J.M.L.: Investigation, Writing- review & editing.

B.E.M.: Investigation, Methodology, Writing- review & editing.

A.N.R.: Investigation, Methodology, Writing- review & editing.

L.A.R.: Conceptualization, Data curation, Funding acquisition, Investigation, Methodology, Project administration, Supervision, Writing- original draft.

