## Supplemental for "Dynamic fates of dietary antigen-specific T helper cells in a model of early life oral tolerance"

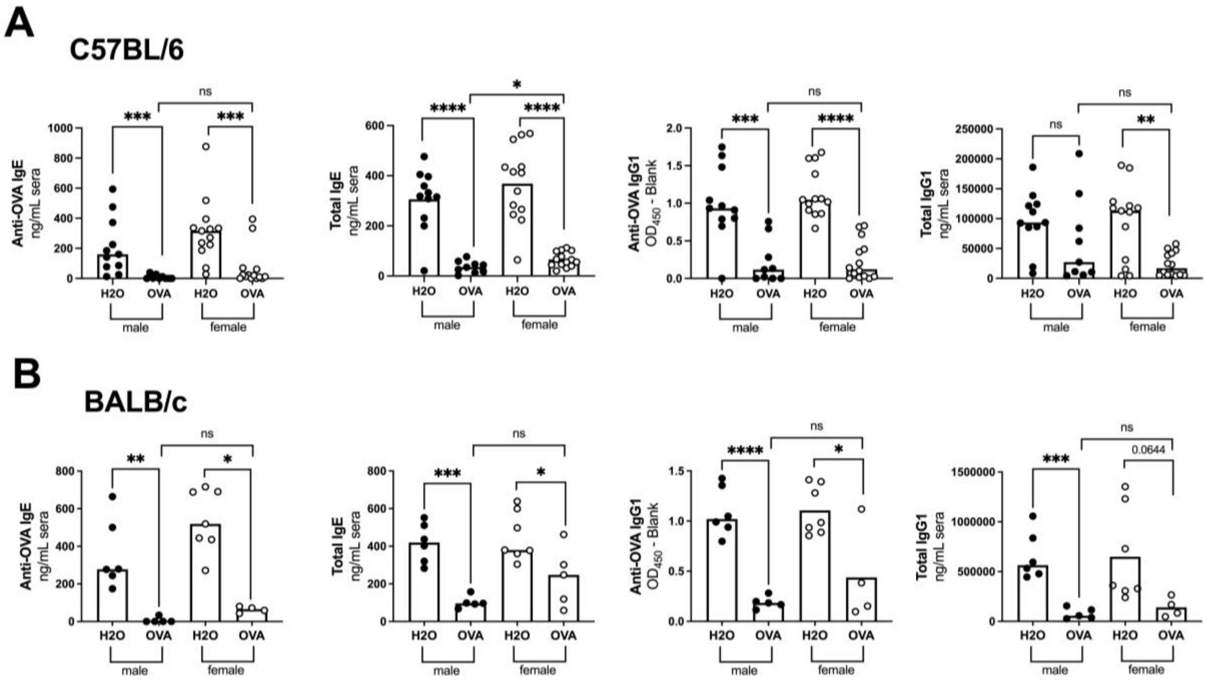


**Supplemental Figure 1. Both female and male C57BL/6 and BALB/c mouse pups develop oral tolerance to 2mg daily oral OVA.** The same data shown in Figure 1 were assessed for the effect of sex on OVA-specific antibody levels. Male (●) and female (○) mouse pups were orally gavaged with $H_2O$ or 2mg of ovalbumin (OVA) in $H_2O$ on days 16-20 of life. All mice received systemic intraperitoneal (IP) challenges with 100µg OVA mixed with 1mg aluminum hydroxide (Alum) on days 22 and 36 of life. On day 43 of life blood was collected from each mouse to assess the development of oral tolerance. OVA-specific IgE, total IgE, OVA-specific IgG1 and total IgG1 in C57BL/6 **(A)** and BALB/c mice **(B)** were quantified by ELISAs in the serum. Column heights are at the mean and data were analyzed by an Unpaired t-test for normally distributed data. Column heights are at the median and data were analyzed by a Mann-Whitney test for non-normally distributed data. Data shown are pooled from 3 (A) or 2 (B) independent experiments with 1-6 male or female mice per experimental group in each experiment. * = $p<0.05$, **=$p<0.01$, ***=$p<0.001$, ****=$p<0.0001$, ns = no statistical difference, p values ≤0.1 are presented on graphs.

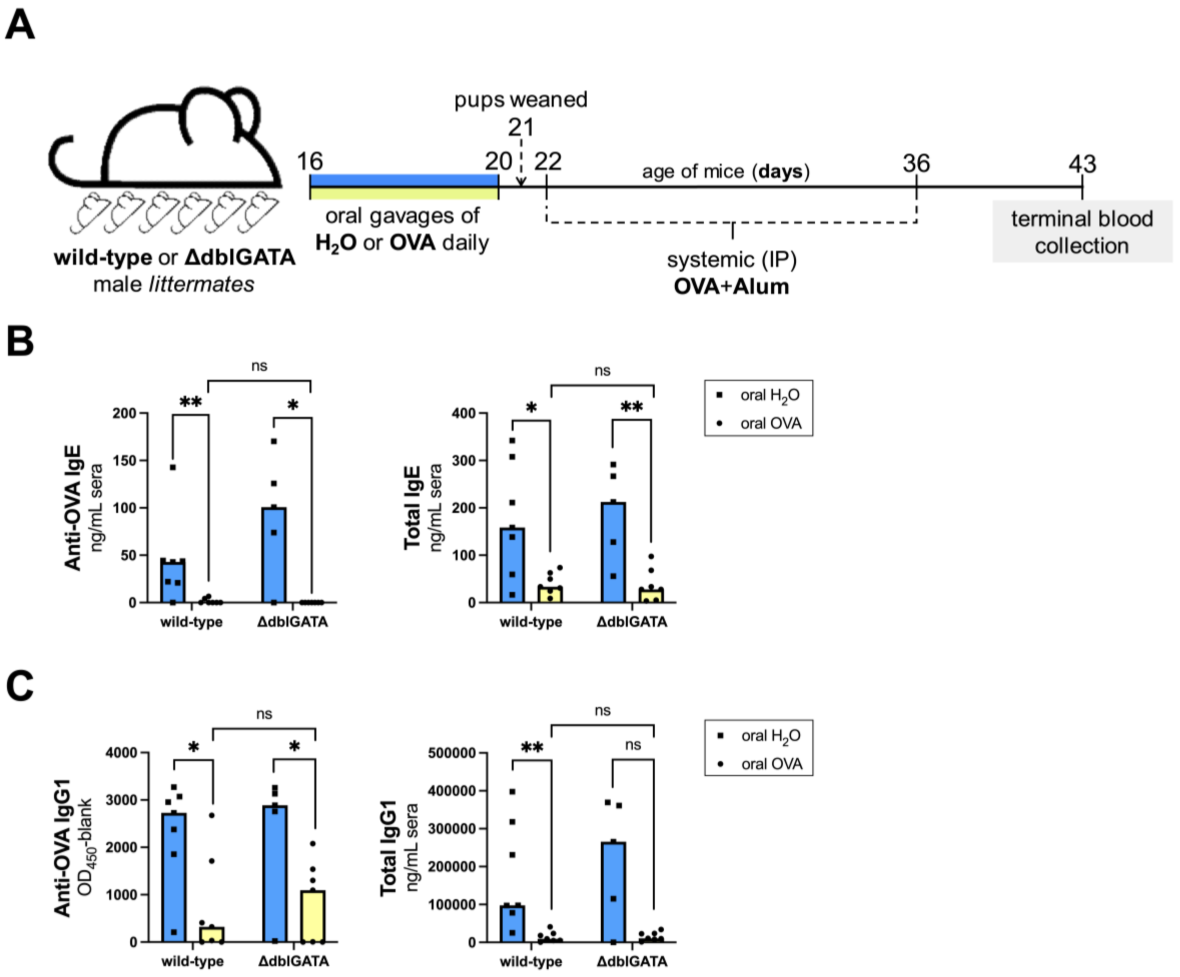


**Supplemental Figure 2. Eosinophils are not essential for the development of oral tolerance in early life. (A)** Experimental timeline: Male wild-type and ΔdblGATA C57BL/6 littermate controls were orally gavaged with $H_2O$ or 2mg of ovalbumin (OVA) in $H_2O$ on days 16-20 of life. All mice received systemic intraperitoneal (IP) challenges with 100µg OVA mixed with 1mg aluminum hydroxide (Alum) on days 22 and 36 of life. On day 43 of life blood was collected from each mouse to assess the development of oral tolerance. **(B)** Serum OVA-specific and total IgE and **(C)** serum OVA-specific and total IgG1 levels were quantified by ELISA. Column heights are at the median and data were analyzed by a Mann-Whitney test. Data shown are pooled from 2 independent experiments with 2-5 mice per experimental group in each experiment. *=$p<0.5$, **=$p<0.01$, ns = no statistical difference.

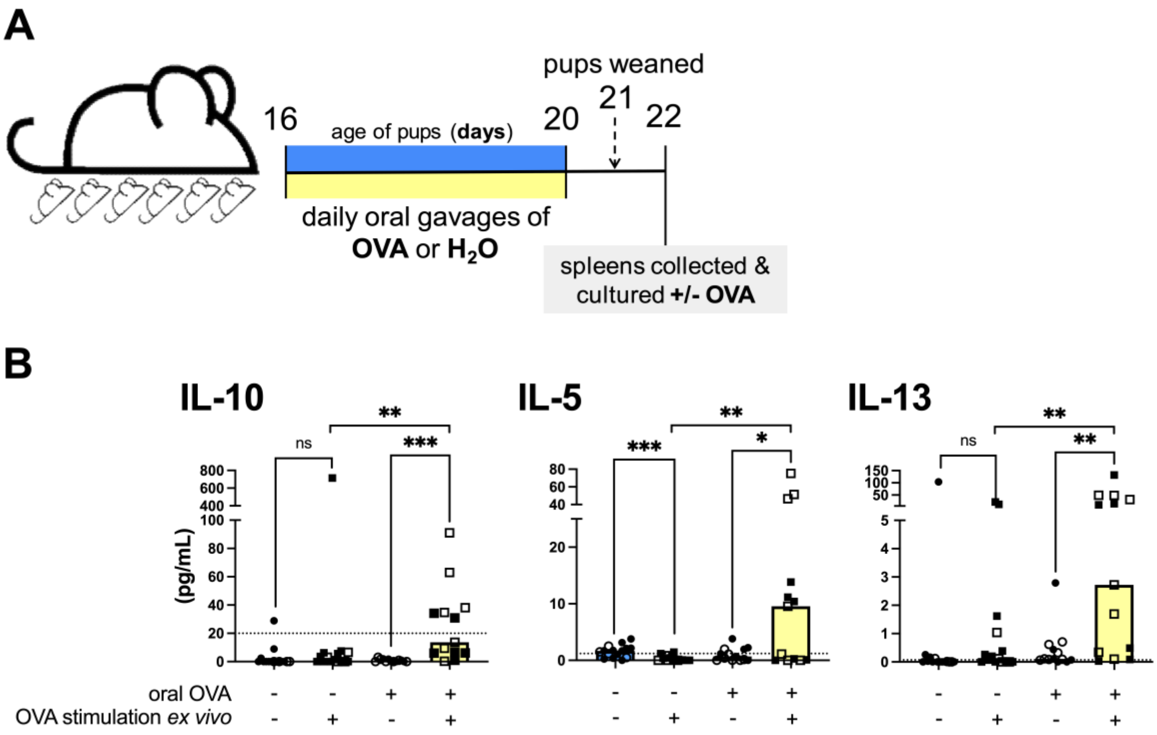


**Supplemental Figure 3. Early life oral OVA exposure leads to the OVA-specific production of Treg and Th2 associated cytokines. (A)** Experimental timeline: male (●, ■) and female (○, □). C57BL/6 mouse pups were orally gavaged with 2mg of OVA in $H_2O$ or $H_2O$ alone on days 16-20 of life. Spleens were collected on day 22 of life and $1x10^6$ splenocytes were cultured in the presence or absence of 10μg/mL OVA for 48hrs. **(B)** IL-10, IL-5 and IL-13 were quantified in culture supernatants using a cytometric bead array assay. Dotted lines indicate the lower limit of accurate quantification. Column heights are at the median and data were analyzed by a Wilcoxon matched-pairs signed rank test for paired experimental groups (unstimulated versus stimulated) and a Mann-Whitney test for unpaired experimental groups (oral $H_2O$ versus oral OVA). Data shown are pooled from 2 independent experiments with 6-9 mice per experimental group in each experiment. *=p<0.5, **=p<0.01, ***=p<0.001, ns = no statistical difference.

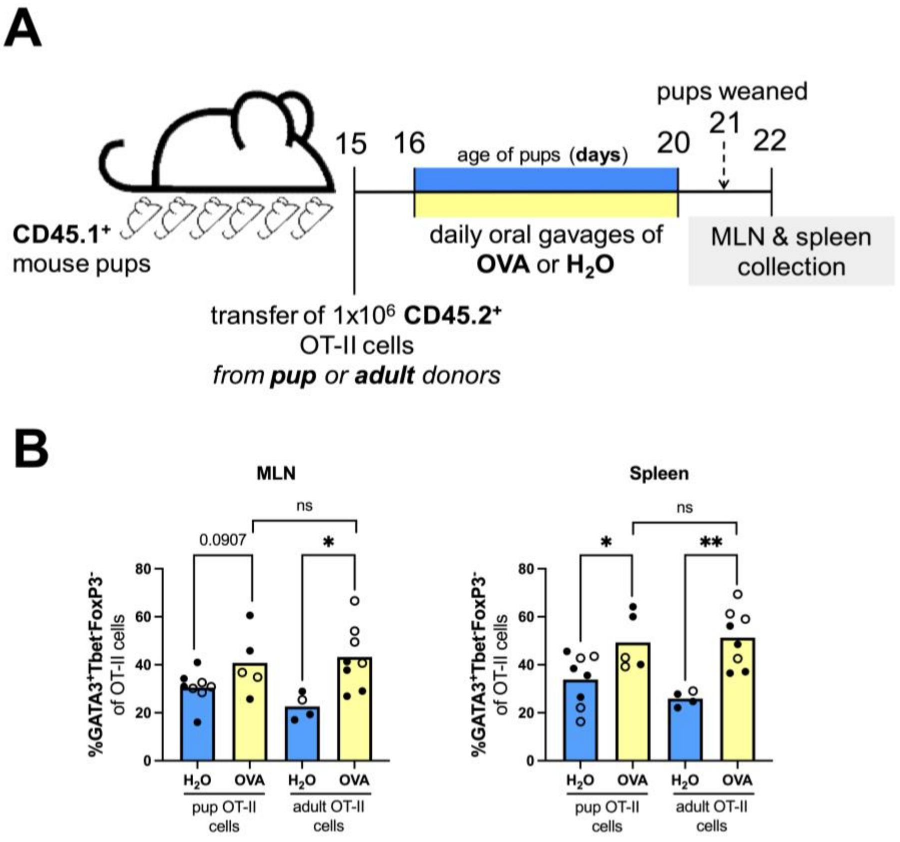


**Supplemental Figure 4. Mouse pup and adult OT-II donor cells develop into OVA-specific Th2 cells following early life oral OVA exposure.** Data from of the independent experimental repeats presented in Figure 5 was originally collected alongside mice who received adult OT-II cells to assess the effect of the age of OT-II cells on Th2 responses. **(A)** Experimental timeline: 15 day old male (●) and female (○) CD45.1 C57BL/6 mouse pups injected with 1x10$^6$ OT-II cells isolated from 3-week-old (pup) or 8-week-old (adult) OT-II donor mice and then orally gavaged with 2mg of ovalbumin (OVA) in $H_2O$ or $H_2O$ alone on days 16-20 of life. On day 22 of life the mesenteric lymph node (MLN)s and spleen were collected for flow cytometry. **(B)** Frequencies of Th2 cells (GATA3$^+$T-bet$^-$FoxP3$^-$) among OT-II cells (CD45.2$^+$CD45.1$^-$CD4$^+$CD3$^+$ live singlets) in the MLNs and spleen. Column heights are at the mean and data were analyzed by an Unpaired t-test. Data shown are from 1 experiment with 4-8 mice per experimental group. *=p<0.5, **=p<0.01, ns = no statistical difference, p values ≤0.1 are presented on graphs.
